# Infralimbic prefrontal cortical projections to the autonomic brainstem: Quantification of inputs to cholinergic and adrenergic/noradrenergic nuclei

**DOI:** 10.1101/2025.04.17.649459

**Authors:** Ema Lukinic, Tyler Wallace, Carlie McCartney, Brent Myers

## Abstract

The ventromedial prefrontal cortex regulates both emotional and physiological processes. The infralimbic cortex (IL), a prefrontal subregion in rodents, integrates behavioral, neuroendocrine, and autonomic responses to stress. However, the organization of cortical inputs to brainstem nuclei that regulate homeostatic responses are not well defined. We hypothesized that IL projections differentially target pre-ganglionic parasympathetic neurons and adrenergic/noradrenergic nuclei. To quantify IL projections to autonomic brainstem nuclei in male rats, we utilized viral-mediated gene transfer to express yellow fluorescent protein (YFP) in IL glutamatergic neurons. YFP-positive projections to cholinergic and adrenergic/noradrenergic nuclei were then imaged and quantified. Cholinergic neurons were visualized by immunohistochemistry for choline acetyltransferase (ChAT), the enzyme responsible for the synthesis of acetylcholine. Adrenergic/noradrenergic neurons were visualized with immunohistochemistry for dopamine beta-hydroxylase (DBH) which converts dopamine to norepinephrine. Our results indicate that IL glutamate neurons innervated the cholinergic dorsal motor nucleus of the vagus with greater density than the nucleus ambiguus. Furthermore, numerous DBH-positive cell groups received IL inputs. The greatest density was to the C2 and A2 regions of the nucleus of the solitary tract with intermediate levels of input to A6 locus coeruleus and throughout the C1 and A1 regions of the ventrolateral medulla. Minimal input was present in the pontine A5. Additionally, IL projections targeted the local GABAergic neurons that regulate activity within preautonomic nuclei. Collectively, our results indicate that IL pyramidal neurons project to vagal preganglionic parasympathetic neurons, presympathetic neurons of the ventrolateral medulla, as well as diffuse homeostatic modulators the nucleus of the solitary tract and locus coeruleus. Ultimately, these findings provide a roadmap for determining circuit-level mechanisms for neural control of homeostasis and autonomic balance.

## Introduction

The autonomic nervous system maintains physiological homeostasis through the regulation of visceral functions including blood pressure, heart rate, and metabolism. Critically, there is a complex interplay between autonomic physiology and emotional processes that has been hypothesized to underly the co-occurrence of mood disorders and cardiometabolic pathologies (Grippo and Johnson 2009; Steptoe and Kivimäki 2012; Sgoifo et al. 2015; Wulsin et al. 2018). Thus, identifying neurobiological pathways that integrate cognitive and emotional processes with systemic function is essential to understand the brain basis of health. The ventromedial prefrontal cortex (vmPFC) regulates both affective processes and physiological functions (Damasio 1996; Gianaros and Sheu 2009; Thayer et al. 2012; Wallace and Myers 2021). Specifically, Brodmann’s area 25 (BA25), a subregion of the vmPFC, has been linked to mood disorders (Drevets et al. 1997, 1998) and identified as a component of the central autonomic network (Kimmerly et al. 2005; Beissner et al. 2013; Shoemaker et al. 2015). Human imaging studies link BA25 with the processing of emotional and social stimuli; moreover, multiple psychiatric conditions relate to altered structure and function of BA25 (Mayberg et al. 1999, 2005). Notably, BA25 also modulates cardiovascular functions, including heart rate and blood pressure (Ziegler et al. 2009; Lacuey et al. 2018). However, the organization of vmPFC projections to autonomic motor outputs have not been well defined.

The infralimbic cortex (IL) is a subregion of rodent prefrontal cortex with homologous cytoarchitecture and connectivity to BA25 (Öngür and Price 2000). Accordingly, the IL has been investigated as a regulator of emotional behaviors including fear, motivation, sociability, and stress coping (Myers-Schulz and Koenigs 2012). Furthermore, pharmacological manipulation of IL activity alters heart rate and blood pressure responses to acute stressors (Resstel and Corrêa 2006; Tavares and Corrêa 2006; Resstel et al. 2006; Tavares et al. 2009). Genetic approaches to limit output specifically from IL pyramidal glutamate neurons increase endocrine and cardiovascular responses to stress (Myers et al. 2017a; Schaeuble et al. 2019). Further, optogenetic activation of male IL glutamatergic neurons increases social motivation coupled with reductions in blood pressure, heart rate, and glucoregulatory responses to acute stress (Wallace et al. 2021). Thus, IL glutamatergic neural activity is both necessary and sufficient to integrate organismal responses to environmental challenge. Prior investigations of IL outflow have focused on projections to forebrain nuclei (Hurley et al. 1991; Vertes 2004), which are hypothesized to relay cortical information to preautonomic nuclei (Ulrich-Lai and Herman 2009). The IL densely innervates numerous forebrain nuclei involved in homeostatic regulation, including specific regions of the hypothalamus, amygdala, and bed nucleus of the stria terminalis (Vertes 2004; Myers et al. 2014; Wood et al. 2019). Although specific prefrontal projections to the forebrain modulate key behavioral aspects of adaptation (Radley et al. 2009; Warden et al. 2012; Adhikari et al. 2015; Johnson et al. 2019), cortical projections to pre-autonomic brainstem nuclei may mediate contextual and emotional influences on physiological integration.

Prior analyses of IL anterograde projections that examined the brainstem used the tracer *Phaseolus vulgaris*-leucoagglutinin (Pha-L) to qualitatively assess innervation patterns and yielded differing results. For instance, Hurley et al. (Hurley et al. 1991) reported that IL efferents target multiple autonomic-related nuclei throughout the hindbrain including the dorsal vagal complex and ventral medulla. However, Vertes (Vertes 2004) found IL projections to the periaqueductal gray (PAG) with minimal innervation of the ventral hindbrain. Moreover, dextran amine tracing from the IL combined with ultrastructural microscopy indicated that IL efferents made synapses onto tyrosine hydroxylase (TH)-positive neurons in the rostral ventrolateral medulla (RVLM) and nucleus of the solitary tract (NTS), as well as TH-negative neurons of unknown neurochemistry (Gabbott et al. 2007). Thus, while there is evidence for IL innervation of the autonomic brainstem, the relative density and potential lateralization of these projections, specific subregions targeted, and neurochemical specificity of the circuitry have not been quantitatively assessed. To analyze IL projections to the brainstem, we utilized viral-mediated gene transfer to express YFP in IL glutamatergic neurons. Imaging of brainstem nuclei was then guided by immunohistochemical visualization of cells that synthesize epinephrine/norepinephrine, acetylcholine, or GABA to quantify the relative density of YFP-positive glutamatergic projections to preganglionic parasympathetic neurons and pre-sympathetic nuclei. Collectively, the current findings outline a road map for data-driven analysis of cortical-brainstem circuitry.

## Methods

### Animals

Adult male Sprague-Dawley rats were obtained from Envigo (Denver, CO) with weights ranging from 250-300 g. After stereotaxic surgery, rats were housed individually in shoebox cages with cardboard tubes for enrichment in a temperature-and humidity-controlled room with a 12-hour light-dark cycle (lights on at 07:00h, off at 19:00h) and food and water *ad libitum*. All procedures and protocols were approved by the Colorado State University Institutional Animal Care and Use Committee (protocol: 16-6871A) and complied with the National Institutes of Health Guidelines for the Care and Use of Laboratory Animals. Signs of poor health and/or weight loss ≥ 20% of pre-surgical weight were *a priori* exclusion criteria. These criteria were not met by any animals in the current experiments.

### Stereotaxic surgery

For microinjections, rats were anesthetized with isoflurane (1-5%) followed by analgesic (0.6 mg/kg buprenorphine-SR, subcutaneous) administration. Rats received unilateral microinjections (50 nL) of adeno-associated virus (AAV) into the IL (2.7 mm anterior to bregma, 0.6 mm lateral to midline, and 4.2 mm ventral from dura). The AAV5-packaged construct (University of North Carolina Vector Core, Chapel Hill, NC) expressed yellow fluorescent protein (YFP) under the CaMKIIα promoter for pyramidal cell-predominant expression (Wood et al. 2019). All microinjections were carried out with a 25-gauge, 2-µL microsyringe (Hamilton, Reno, NV) using a microinjection unit (Kopf, Tujunga, CA) at a rate of 5 minutes/µL. The needle was left in place for 5 minutes before and after injections to reduce tissue damage and allow diffusion. Skin was closed with wound clips that were removed 2 weeks after surgery. All animals were allowed 6 weeks for YFP expression.

### Tissue collection

Rats were injected with sodium pentobarbital (100 mg/kg) and perfused transcardially with 0.9% saline followed by 4.0% paraformaldehyde in 0.1 M phosphate buffered saline (PBS). Brains were removed and post-fixed in 4.0% paraformaldehyde for 24 h at room temperature, followed by storage in 30% sucrose in PBS at 4 °C. Coronal sections were made on a freezing microtome (Thermo Fisher Scientific, Waltham, MA) at 30 μm thickness, with a subgroup sectioned sagittally, and stored in cryoprotectant solution at -20 °C until processing.

### Immunohistochemistry

Initial tissue analysis was carried out to identify cases with IL-specific injections. Brain sections were washed in PBS (5 x 5 min) and placed into a 4’,6-diamidino-2-phenylindole (DAPI) solution (300 nM in PBS, RRID: AB_2307445, Thermo Fisher Scientific, Waltham, MA) for 10 min. After another PBS wash (5 x 5 min), the tissue was mounted in polyvinyl medium and cover slipped for imaging.

For labeling choline acetyltransferase (ChAT), sections were first rinsed in PBS (5 x 5 min) at room temperature. Sections were then placed in blocking solution (PBS, 0.1% bovine serum albumin, and 0.2% Triton X-100) for 1 hour. Following blocking, sections were incubated overnight at room temperature in rabbit anti-ChAT primary antibody (1:1000 in blocking solution, RRID: AB_178850, Abcam, Cambridge, UK). Sections were then rinsed in PBS (5 x 5 min) followed by a 1 h incubation in goat anti-rabbit Cy3 secondary antibody (1:500 in PBS, RRID: AB_2307443, Jackson ImmunoResearch, West Grove, PA). The tissue was then washed (5 x 5 min in PBS), mounted with polyvinyl medium, and cover slipped for imaging.

Labeling of dopamine beta-hydroxylase (DBH)-positive neurons was conducted as described above with the following exceptions. Tissue was incubated overnight in mouse anti-DBH primary antibody (1:2500 in blocking solution, RRID: AB_94983, MilliporeSigma, Burlington, MA). Tissue was then incubated for 1 h in donkey anti-mouse Cy3 secondary antibody (1:500 in PBS, RRID: AB_2340813, Jackson ImmunoResearch).

To examine whether IL projections targeted GABAergic neurons within presympathetic cell groups, tissue was immunolabeled for GABA. After a PBS wash, tissue was incubated for 4 h in blocking solution (8% normal goat serum, 6% BSA, 2% normal donkey serum, and 0.4% Triton X-100 in PBS). Sections were then incubated in rabbit anti-GABA primary antibody (1:250 in blocking solution, RRID: AB_11214017; MilliporeSigma) for 60 h. After, the tissue was washed in PBS (5 x 5) and incubated in biotinylated secondary goat anti-rabbit antibody (1:500 in PBS, RRID: AB_2313606; Vector Laboratories) followed by Vectastain ABC Solution for 1 h (1:500 in PBS, RRID: AB_2336827; Vector Laboratories), then Cy3-conjugated streptavidin for 1 h (1:500 in PBS, RRID: AB_2337244; Jackson ImmunoResearch). Finally, the tissue was washed in PBS (5 x 5 min), mounted in polyvinyl medium, and cover slipped for imaging.

### Microscopy, quantification, and analysis

To determine IL injection placement and brainstem anatomy, YFP was imaged with a Zeiss Axio Imager Z2 microscope (Carl Zeiss Microscopy, Jena, Germany) using a 10x objective. The expression of ChAT and DBH aided identification of targeted regions. Images for quantification of YFP density were captured with apotome-enabled optical sectioning (40x objective) in maximum intensity projection renderings from individual z-planes (0.5 μm thickness). Images were collected along the rostral-caudal gradient of each structure, typically representing a rostral, middle, and caudal portion of each nucleus. The Swanson atlas (Swanson 2004) was used to guide anatomical identification with the Paxinos and Watson atlas (Paxinos et al. 2006) used to supplement the identification of catecholamine cell groups. Based on the length of a given region, as few as 2 or as many as 5 image stacks were collected ipsilateral to each injection site. To investigate lateralization, regions were also imaged contralateral to the injection site. Due to significant autofluorescence in the brainstem, all quantitative imaging of the YFP channel was accompanied by imaging in the Cy5 channel as a measure of nonspecific signal. Here, YFP relative density was corrected with Cy5 signal to increase the specificity of quantification.

To quantify the regional density of IL input, optical projections were processed individually in ImageJ Fiji (ver 1.51N). Following the division of the YFP signal by the Cy5 channel, thresholding was applied manually and the percent area occupied by fluorescence was quantified by a blinded observer. For regions of interest, the mean percent area from each location along the rostral-caudal gradient was averaged to yield a regional percent area value in each of the 3 cases. These values were then corrected for the case median percent area of all regions sampled to account for between-case variation in construct expression. The regional median-corrected percent area of YFP density was then averaged across all 3 cases. All data are expressed as mean ± standard error of the mean (SEM). Data were visualized with Prism 9.3.1 (GraphPad, San Diego, CA).

## Results

### Injection placement and quantitative approach

Following exclusion of cases with spread to PL (n = 2), 3 cases with IL-specific injections (Fig. 1A) were used for quantitative analysis. Cases were injected unilaterally in the IL with side randomized. The final analysis included 2 left and 1 right. Injections were primarily targeted to the ventral aspects of the deep layers of the mid IL, with spread to more superficial layers predominately in rostral areas of the IL. YFP expression saturated much of the rostral and mid IL, with limited expression in the caudal IL and an absence of dorsal spread into the PL. An additional 2 IL-specific injections were used for sagittal examination of major fiber tracts into the brainstem (Fig. 1B). Sagittal examination pointed to gradients in the density of YFP-expressing projections. Generally, YFP was robustly expressed throughout the midbrain encompassing dorsal areas near the PAG. As projections traveled caudally toward the medulla, parallel tracts emerged with high density of innervation in the dorsal vagal complex, as well as a projection through the ventral medulla. More sparse labeling of projections continued to the spinal cord. To quantify projection density, YFP was imaged in high-magnification z-stacks throughout regions of interest. To enhance signal specificity and remove autofluorescence signals from analysis, Cy5 was imaged and the YFP signal was corrected for Cy5 fluorescence, yielding a specific measure of IL glutamate projection density (Fig. 1C). As described in the methodology, the % density of YFP in each region of interest was corrected for the median % area for each case; thus, reported projection density was normalized across cases.

**Figure 1.**
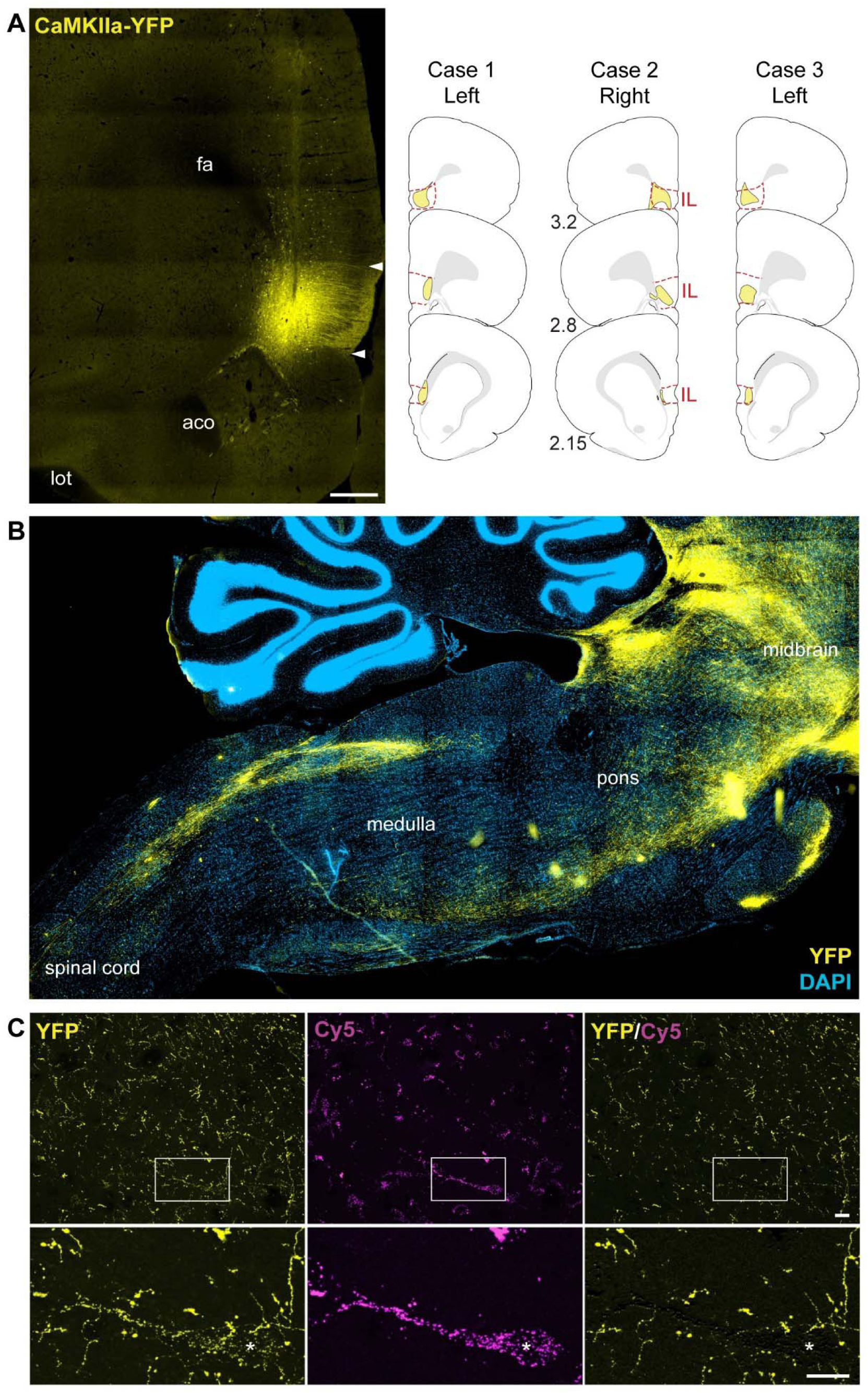
A: Representative photomicrograph of injection in IL (left) and injection maps for each case (right) with schematics adapted from Swanson (Swanson 2004). Distances are anterior to bregma, scale bar: 500 μm. B: Sagittal view of YFP-expressing IL projections. C: Representative approach to quantification from the ventral medulla. In all regions, YFP was imaged followed by Cy5 correction to account for autofluorescence or other non-specific signals. White boxes indicate area of higher magnification insets (bottom). Scale bars: 10 μm, * indicates the location of corrected autofluorescent cell body. aco: anterior commissure, CaMKIIa: calcium–calmodulin-dependent protein kinase II alpha, DAPI: 4’,6-diamidino-2-phenylindole, fa: anterior forceps of the corpus callosum, IL: infralimbic cortex, lot: lateral olfactory tract, YFP: yellow fluorescent protein.

### Projections to cholinergic preganglionic parasympathetic neurons

Coronal sections with ChAT immunoreactivity were imaged throughout the dorsal vagal complex to identify the relative distribution of YFP-expressing IL projections to the dorsal motor nucleus of the tenth cranial nerve (DMX; Fig 2A). The DMX gives rise to a large portion of the preganglionic parasympathetic neurons that generate the vagus nerve (Espinoza et al. 2021). Higher magnification micrographs were imaged with Cy5 correction for quantification of IL projections. Similar approaches were employed to investigate the nucleus ambiguus (NAmb: Fig 2B) which represents a population of preganglionic parasympathetic neurons accounting for the majority of cardiac parasympathetic regulation (Espinoza et al. 2021). Quantitative results found that IL glutamate neuron projections provided minimal input to NAmb with greater density of inputs to the DMX bilaterally (Fig 2C).

**Figure 2.**
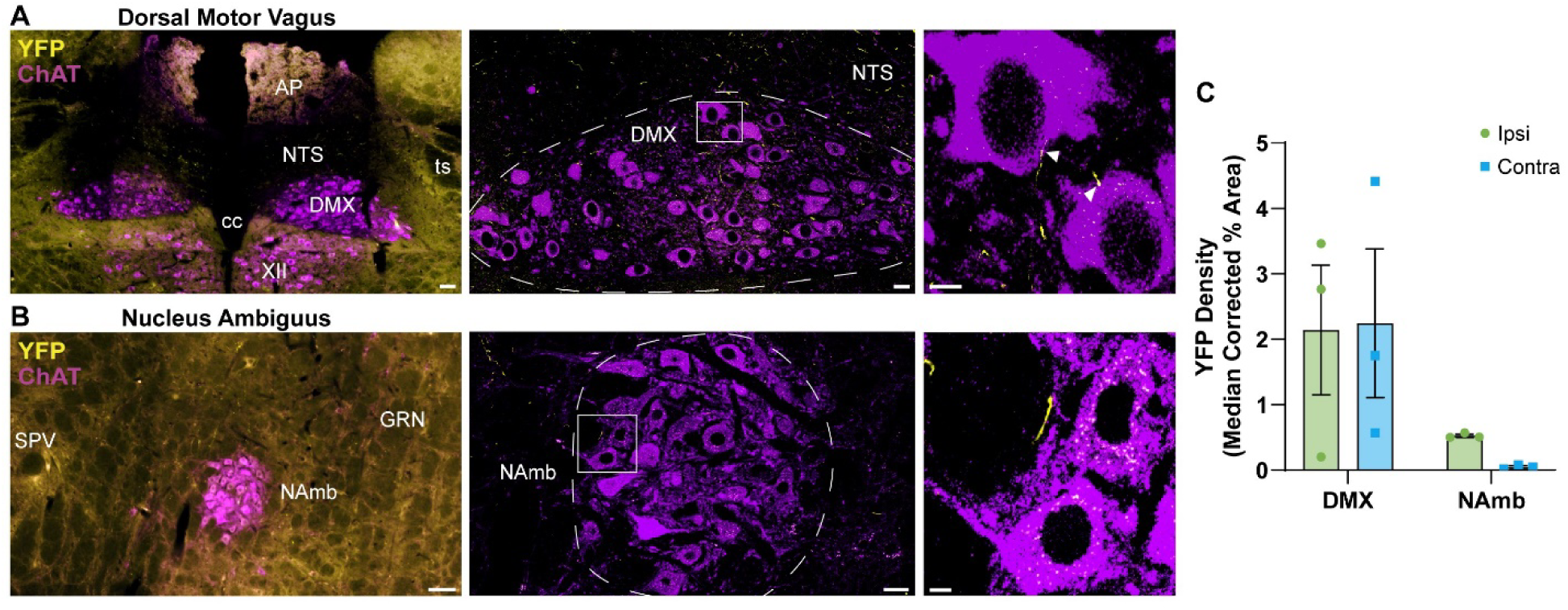
A: Tiled images were used to identify the location of cholinergic preganglionic neurons in the DMX (left), with high-magnification z-stacks corrected for Cy5 fluorescence utilized in quantitative analysis of IL projection density (middle). White boxes highlight the location of insets displaying appositions (white arrows; right). B: Similar approaches were used to examine NAmb. C: Quantification of projection density as a function of laterality with injection sites. Data are presented as mean ± SEM. Scale bars: 100 μm (left), 10 μm (middle), 5 μm (right). XII: cranial nerve 12 hypoglossal, AP: area postrema, cc: central canal, ChAT: choline acetyltransferase, DMX: dorsal motor nucleus of the vagus, GRN: gigantocellular reticular nucleus, NAmb: nucleus ambiguus, NTS: nucleus of the solitary tract, SPV: spinal trigeminal nucleus, ts: solitary tract, YFP: yellow fluorescent protein.

### Projections to norepinephrine/epinephrine-producing neurons

DBH immunoreactivity guided imaging of the primary norepinephrine (A) and epinephrine (C) cell groups. These pre-sympathetic nuclei target spinal preganglionic sympathetic neurons, as well as ascending forebrain circuits. Indeed, the locus coeruleus (A6, Fig 3A), NTS (C2 and A2, Fig 3B), and VLM (C1 and A1, Fig 3C) provide most of the epinephrine and norepinephrine to the central nervous system (Dahlström and Fuxe 1964; Sawchenko and Swanson 1981; Myers et al. 2017b). Analysis also included A5 which yielded almost no YFP expression (Fig 3D). Overall, the highest density of IL input was found in the C2 region of the NTS ipsilaterally and the A2 bilaterally. There was also moderate IL projection density ipsilaterally throughout the C1 RVLM, A1 caudal VLM (CVLM), and A6 LC.

**Figure 3.**
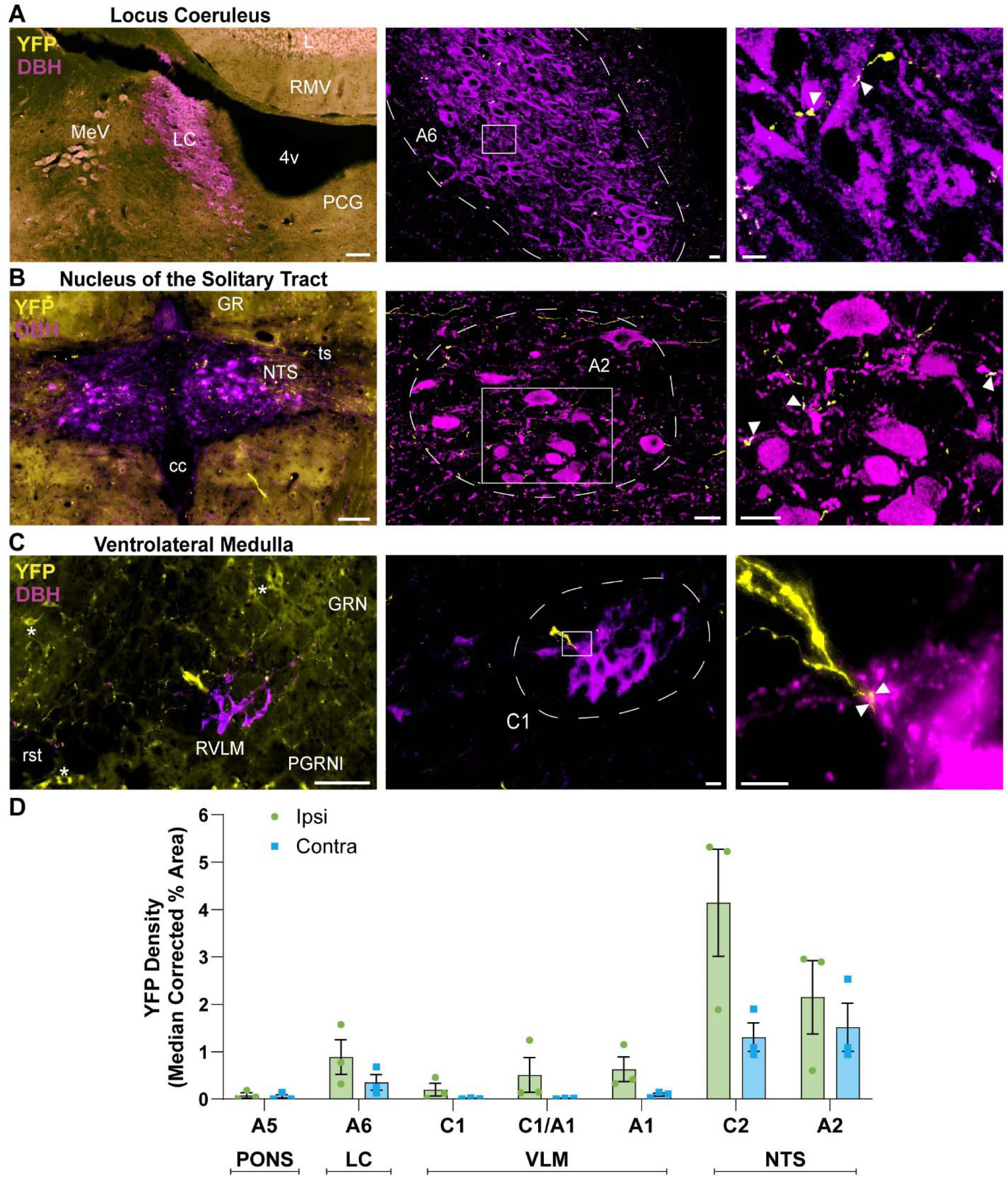
A: Tiled images were used to identify the location of catecholaminergic neurons in the LC (left), with high-magnification z-stacks corrected for Cy5 fluorescence utilized in quantitative analysis of IL projection density (middle). White boxes highlight the location of insets displaying appositions (white arrows; right). B: Similar approaches were used to examine NTS and VLM (C). * represent autofluorescent cell bodies removed by subsequent correction. D: Quantification of projection density as a function of laterality with injection sites. Data are presented as mean ± SEM. Scale bars: 50 μm (left), 10 μm (middle), 10 μm (right). 4v: fourth ventricle, cc: central canal, DAPI: 4’,6-diamidino-2-phenylindole, DBH: dopamine beta-hydroxylase, GR: gracile complex, GRN: gigantocellular reticular nucleus, L: lingula of cerebellum, LC: locus coeruleus, MeV: mesencephalic trigeminal nucleus, NTS: nucleus of the solitary tract, PCG: pontine central gray, PGRNl: paragigantocellular reticular nucleus lateral, RMV: rostral medullary velum, rst: rubrospinal tract, ts: solitary tract, VLM: ventrolateral medulla, YFP: yellow fluorescent protein.

### Projections to the periaqueductal gray

Given the density of IL inputs, distinctive morphology, and importance in homeostatic integration, we also examined IL projections throughout the columns of the PAG. As for other nuclei, low magnification YFP imaging was done to identify landmarks (Fig 4A), followed by optical sectioning with correction for non-specific signal (Fig 4B-D). From dorsal to ventral, the dorsal medial (dmPAG; Fig 4B), dorsal lateral (dlPAG; Fig 4C), lateral (lPAG; Fig 4D), and ventral lateral (vlPAG; Fig 4E) regions of the PAG were quantified. The density of IL inputs increased across this dorsal-ventral gradient with midline dmPAG having the least YFP and ipsilateral vlPAG the most (Fig 4F).

**Figure 4.**
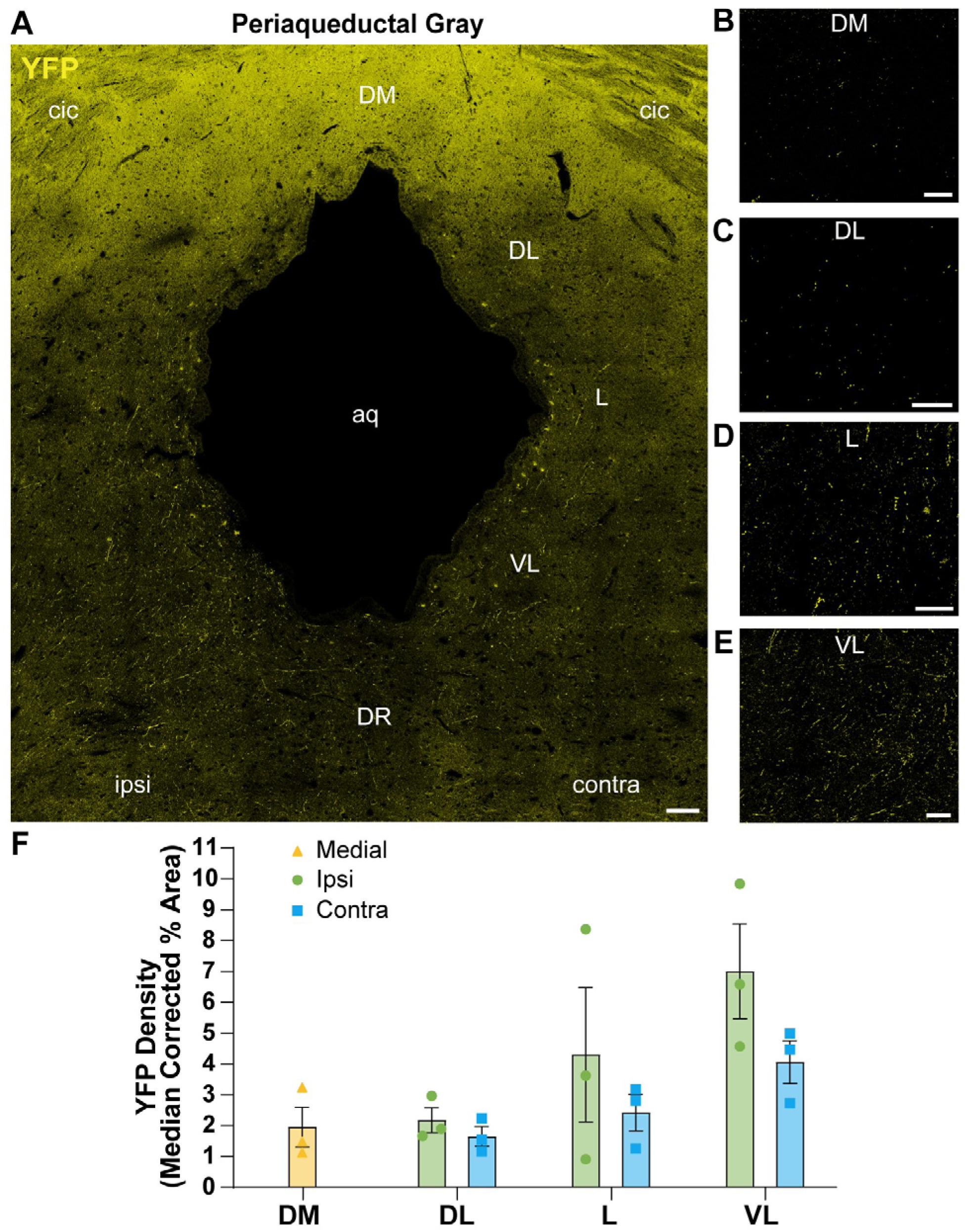
A: Columnar organization of the PAG was imaged bilaterally. Scale bar: 100 μm. Inset images indicate the relative projection density ipsilaterally for the dm (B), dl (C), l (D), and VL (E). Scale bars: 20 μm. F: Quantification of projection density as a function of laterality with injection sites. Data are presented as mean ± SEM. aq: aqueduct, cic: commissure of inferior colliculus, dlPAG: dorsal lateral periaqueductal gray, dmPAG: dorsal medial periaqueductal gray, DR: dorsal raphe, lPAG: lateral periaqueductal gray, vlPAG: ventral lateral periaqueductal gray, YFP: yellow fluorescent protein.

### Projections to GABAergic cells

The IL can inhibit stress responses (Myers et al. 2017a; Schaeuble et al. 2019; Pace et al. 2024), particularly in males (Wallace et al. 2021; Schaeuble et al. 2024), and may be important for context-dependent adaptive processes. Accordingly, we examined whether IL inputs targeted local GABAergic cells in the pre-sympathetic regions that facilitate vigilance behaviors (McCall et al. 2015; Vaaga et al. 2020), fight-or-flight responses (Guyenet et al. 2013), and autonomic balance (Rinaman 2011; Holt 2022). Here, there was IL input to GABA-immunoreactive cells in the PAG (Fig 5A), as well as the GABAergic cells surrounding catecholamine-synthesizing neurons in the LC (Fig 5B), NTS (Fig 5C), and VLM, (Fig 5D). Altogether, indicating that IL post-synaptic targeting is neurochemically diverse and identifying potential pathways for cortical glutamate-mediated inhibition of catecholamine outflow to the central nervous system.

**Figure 5.**
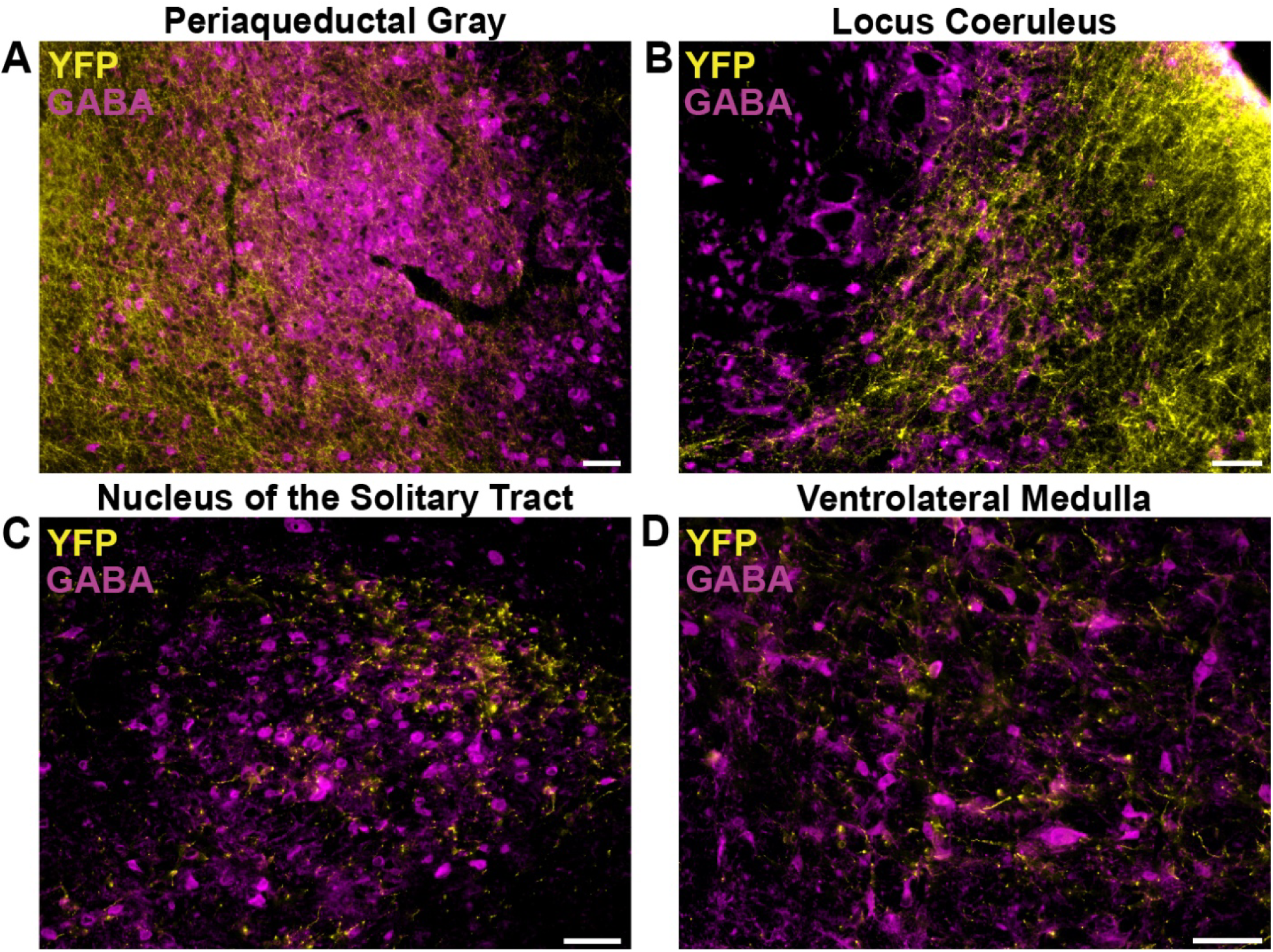
YFP-expressing IL projections targeted GABA-immunoreactive cells in the A: vlPAG, B: peri-LC, C: NTS, D: VLM. Scale bars: 50 μm. IL: infralimbic cortex, LC: locus coeruleus, NTS: nucleus of the solitary tract, PAG: periaqueductal gray, VLM: ventrolateral medulla, YFP: yellow fluorescent protein.

## Discussion

The current study identified direct projections from IL pyramidal neurons to neurochemically-defined brainstem autonomic nuclei. Viral-mediated gene transfer was used to visualize and quantify the density of IL projections, which identified IL glutamate inputs to both catecholaminergic nuclei and preganglionic parasympathetic neurons (Fig 6). By quantifying relative projection density, our data indicate that all cases had extensive IL input to cholinergic vagal motor neurons. Moreover, multiple epinephrine- and norepinephrine-synthesizing cell groups received input from the IL. The highest density was found in the C2 and A2 regions of the NTS with moderate input to the LC and throughout the C1 and A1 regions of the VLM. Additionally, GABA neurons neighboring the catecholamine neurons of the NTS, LC, and VLM also received IL inputs. Altogether, the data suggest that the IL is positioned to directly engage autonomic regulation in a context-specific manner by providing inputs to preganglionic parasympathetic neurons, as well as both sympatho-excitatory and sympatho-inhibitory cell populations. This neurochemically- and anatomically-diverse circuit organization may contribute to the dynamic modulation of organismal physiology based on cognitive and emotional factors.

**Figure 6.**
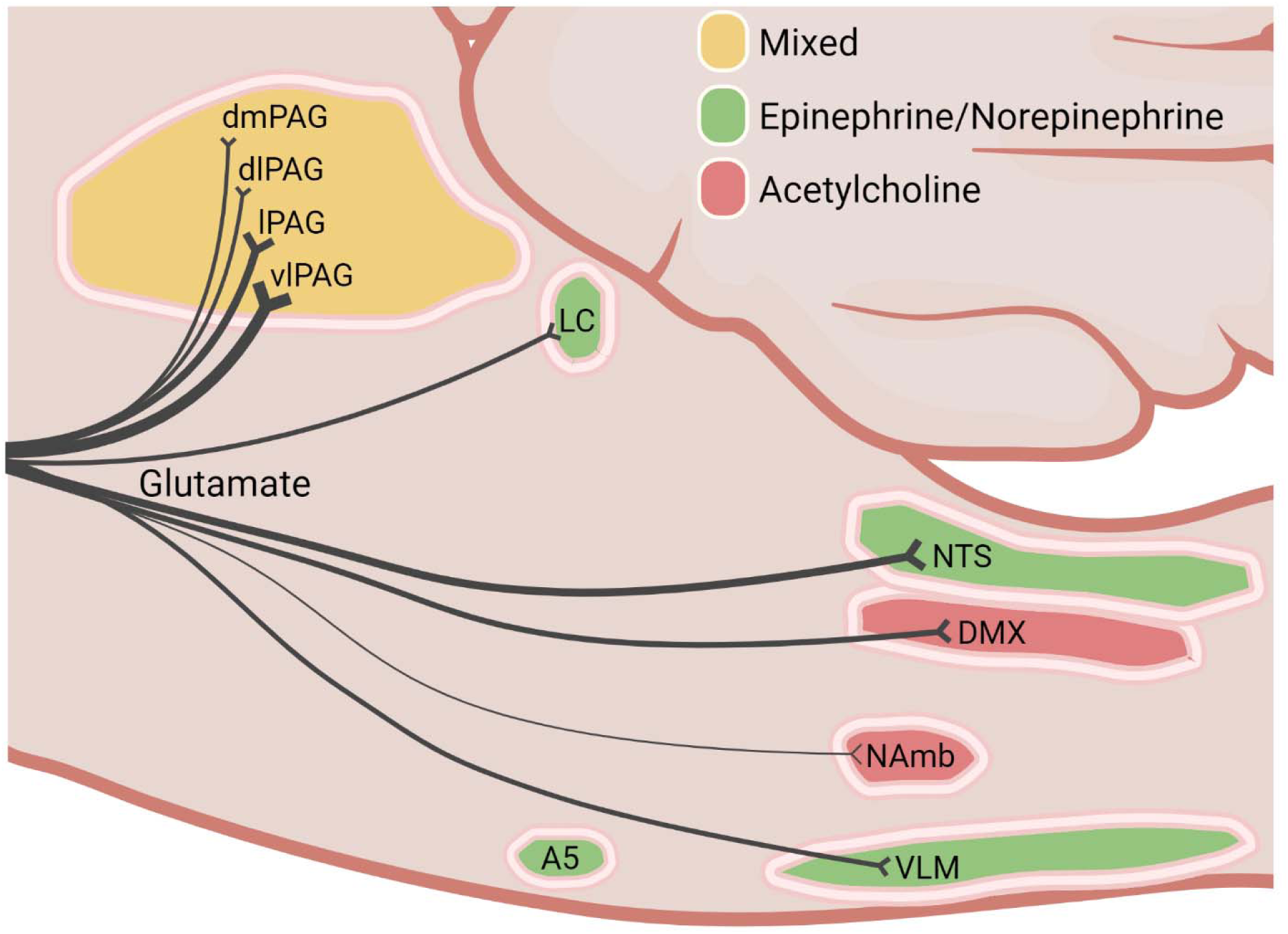
Summary of results: black line thickness represents relative density of IL input to cholinergic nuclei (red), noradrenergic nuclei (green), and the PAG (yellow). The most densely innervated region was the PAG (vl) followed by the NTS, DMX, VLM, and LC. Little or no connectivity was observed in the NAmb and A5. DMX: dorsal motor nucleus of the vagus, IL: infralimbic cortex, LC: locus coeruleus, NAmb: nucleus ambiguus, NTS: nucleus of the solitary tract, VLM: ventrolateral medulla, vlPAG: ventral lateral periaqueductal gray.

The dorsal vagal complex integrates visceral afferent information to fine-tune autonomic motor control (Davis et al. 2004; Browning and Travagli 2011; Herman 2017). The majority of visceral motor regulation is mediated by the cells of the DMX that give rise to the vagus nerve (Browning et al. 2014; Espinoza et al. 2021). These preganglionic parasympathetic neurons innervate postganglionic parasympathetic neurons in the thoracic and abdominal viscera (Ulrich-Lai and Herman 2009). However, the preganglionic parasympathetic neurons that innervate cardiac postganglionic neurons largely originate in the NAmb (Myers 2017; Espinoza et al. 2021). The neurovisceral integration hypothesis was developed to explain the influence of attentional, emotional, and cognitive processes on symaptho-vagal balance and health-related physiology (Thayer and Brosschot 2005; Thayer et al. 2012; Wulsin et al. 2018). Human studies supporting the hypothesis report that impaired prefrontal function decreases the parasympathetic component of autonomic balance (Thayer 2021). However, no studies to our knowledge have examined whether prefrontal neurons directly target preganglionic parasympathetic neurons. The results of the current study indicate that IL glutamate neurons provide a circuit for prefrontal connectivity with preganglionic parasympathetic neurons and suggest relative specificity of these circuits. Particularly, IL neurons target DMX cholinergic neurons to a greater degree than NAmb cholinergic neurons, indicating the capacity for target organ-specific control of parasympathetic outflow. Although, the function of this circuitry remains to be determined.

The NTS is also a component of the dorsal vagal complex and functions as a hub of interoceptive integration (Herman 2017; Kim et al. 2020; Holt and Rinaman 2022; Holt 2022; Baumer-Harrison et al. 2024a). The NTS has varied neurochemistry and synaptic connectivity, influencing a wide array of homeostatic functions (Rinaman 2010; Ghosal et al. 2013; Myers et al. 2017b; Neyens et al. 2020; Baumer-Harrison et al. 2024b; Winzenried et al. 2025). However, the C2 epinephrine-producing and A2 norepinephrine-producing catecholamine neurons have received considerable attention as coordinators of multi-system homeostatic integration (Rinaman 2011; Daubert et al. 2012; Bundzikova-Osacka et al. 2015; Pace and Myers 2024; Carrasco and Breen 2025). The current findings indicate that NTS C2 and A2 neurons are prominent hindbrain targets of IL projections. Moreover, NTS GABAergic neurons also receive IL inputs. Interestingly, NTS GABA neurons have been implicated in hypertension (Sved and Sved 1990; Mohammed et al. 2022), possibly through local actions in the NTS (Bailey et al. 2008), stimulation of vasopressin release (Catelli et al. 1987), and/or inhibition of the DMX (Xu et al. 2015).

The VLM is composed of epinephrine-synthesizing neurons in the C1 RVLM, norepinephrine-synthesizing neurons in the A1 CVLM, and an intermediate zone of mixed C1/A1 cells (Ritter et al. 2019). The VLM generates inputs to spinal preganglionic sympathetic neurons as well as ascending pathways to the hypothalamus (Sawchenko and Swanson 1982; Card et al. 2006). Functional evidence suggests that C1 neurons mediate activation of the sympathetic nervous system (Guyenet et al. 2001, 2013; Stornetta and Guyenet 2018; Zsombok et al. 2024), while A1 neurons stimulate stress hormone release via the hypothalamic-pituitary-adrenal axis (Li et al. 2018). Thus, the VLM network integrates numerous neural inputs to control sympathetic motor responses and endocrine physiology (Schreihofer and Guyenet 2002; Madden and Sved 2003; Desmoulins et al. 2025). The current results indicate that IL projections broadly innervate the multiple subregions of the VLM, in addition to the GABAergic neurons that regulate local network activity. Importantly, GABAergic and glycinergic neurons in the CVLM provide inhibitory signaling to the RVLM to mediate the baroreflex (Schreihofer and Guyenet 2002; Mandel and Schreihofer 2008; Gao et al. 2019). Recent studies of IL synapses in the RVLM found that circuit stimulation reduces HPA axis and sympathetic stress responses, possibly through engagement of GABAergic cells to limit catecholaminergic outflow (Pace et al. 2024).

The LC houses norepinephrine-expressing neurons that broadly innervate the central nervous system (Dahlström and Fuxe 1964; Swanson 1976; Schwarz and Luo 2015). The LC is the primary source of norepinephrine for the cortex and hippocampus and modulates behaviors related to arousal, attention, and stress responses (Aston-Jones et al. 1999; Francis et al. 1999; Valentino and Van Bockstaele 2008; Wood et al. 2017). The current studies found direct IL inputs to the LC as well as the GABAergic cells in the peri-LC region. The GABA neurons adjacent to LC norepinephrine neurons act as interneurons to regulate LC function (Aston-Jones et al. 2004; Breton-Provencher and Sur 2019). Thus, this diverse post-synaptic targeting may allow the IL to either facilitate or gate central norepinephrine activity depending on context. The midbrain PAG regulates numerous adaptive functions, including analgesia, autonomic responses, and behaviors related to defense, fear, and anxiety (Bandler and Shipley 1994; Behbehani 1995). The columnar organization of the PAG generates distinct connectivity and individualized functions with discrete behavioral responses initiated by neighboring columns of the PAG. For instance, the dmPAG and dlPAG mediate active defensive behaviors such as flight, while the lPAG and vlPAG facilitate immobile defensive strategies including freezing (Assareh et al. 2016; La-Vu et al. 2022). These canonical behaviors are thought to be mediated by glutamate neurons and recent examination of pain behaviors found opposing actions for glutamate and GABA neurons in the vlPAG. Here, chemogenetic activation and inhibition approaches indicated that vlPAG glutamate neurons suppress nociception while GABA neurons facilitate nociceptive responses (Samineni et al. 2017). The present analysis identified IL projections to the PAG that increase in density along the dorsal to ventral gradient with vlPAG receiving the most robust input. Moreover, a portion of PAG cells receiving cortical inputs are GABAergic.

The current study corroborates prior reports of IL efferents to the brainstem while making new contributions based on the quantification of projection density and identification of post-synaptic neurochemistry. Early studies found anterograde Pha-L projections to autonomic cell groups including the VLM and dorsal vagal complex, predominantly the NTS (Hurley et al. 1991). However, other studies identified anterograde labeling in the PAG, LC, and NTS with limited inputs to the ventral hindbrain (Vertes 2004). Retrograde tracing from the VLM labeled cells in the vmPFC, primarily the IL (Gabbott et al. 2005) and dextran amine anterograde tracing provided further support for a PFC-VLM pathway (Gabbott et al. 2007). The later studies utilized electron microscopy to visualize IL synaptic contacts with TH-expressing catecholamine neurons in the VLM and NTS, as well other non-catecholaminergic neurons. Data from the current study suggest that these cells may, in part, be GABAergic neurons. The present results also add examination of relative projection density across norepinephrine-producing cells and establish direct IL inputs to vagal preganglionic motor neurons. Furthermore, while innervation of the DMX was bilateral, most other regions received ipsilateral-dominant input from the IL.

There are limitations to the current approaches that should be considered when interpreting outcomes. Primarily, the variable cytoarchitecture of brainstem nuclei limited the ability to specifically quantify expression of presynaptic terminals as in prior forebrain studies (Wood et al. 2019; Pace et al. 2020). Thus, the present report utilized a genetically-encoded YFP construct that labels both axons and synapses. Accordingly, the results may overestimate the presynaptic input across regions of interest.

Additionally, there are numerous brainstem nuclei with a variety of neurotransmitter populations that may be important targets of the IL for orchestrating organismal homeostasis. For instance, the serotonergic raphe nuclei and mixed cell populations of the parabrachial nucleus have been reported to receive IL input (Hurley et al. 1991; Vertes 2004). However, the current investigation focused on preganglionic parasympathetic neurons and major sites of catecholamine regulation, without attempting to identify all brainstem targets of IL. Furthermore, this study only included male rats and emerging evidence suggests considerable differences between male and female neurobiology (Rich-Edwards et al. 2018). Additional examination of female cortical-brainstem connections may yield sex differences.

Overall, a genetically-encoded reporter of IL glutamate projections combined with visualization of neurochemically-defined cell populations identified relative patterns of cortical input across the brainstem. These circuit connections may be critical for homeostatic adaptation and open the door for data-driven functional studies to query region-specific circuit and synaptic function.

## Acknowledgments

AAV5-packaged vectors for CaMKIIa-YFP were provided by the University of North Carolina Vector Core under material transfer agreement with Karl Deisseroth and Stanford University. This work was supported by R01 HL150559 to B. Myers.

## Acronyms

4v: fourth ventricle
XII: cranial nerve 12, hypoglossal
A2, A5, A6: norepinephrine cell groups
AAV: adeno-associated virus
C1, C2: epinephrine cell groups
aco: anterior commissure
AP: area postrema
aq: aqueduct
CaMKIIa: calcium–calmodulin-dependent protein kinase II alpha
cc: central canal
ChAT: choline acetyltransferase
cic: commissure of inferior colliculus
CVLM: caudal ventrolateral medulla
DAPI: 4’,6-diamidino-2-phenylindole
DBH: dopamine beta-hydroxylase
dlPAG: dorsal lateral periaqueductal gray
dmPAG: dorsal medial periaqueductal gray
DMX: dorsal motor nucleus of the vagus
DR: dorsal raphe
fa: anterior forceps of the corpus callosum
GR: gracile complex
GRN: gigantocellular reticular nucleus
IL: infralimbic cortex
L: lingula of cerebellum
LC: locus coeruleus
lPAG: lateral periaqueductal gray
lot: lateral olfactory tract
MeV: mesencephalic trigeminal nucleus
NAmb: nucleus ambiguus
NTS: nucleus of the solitary tract
PAG: periaqueductal gray
PCG: pontine central gray
PGRNl: paragigantocellular reticular nucleus, lateral
Pha-L: *Phaseolus vulgaris*-leucoagglutinin
RMV: rostral medullary velum
rst: rubrospinal tract
RVLM: rostral ventrolateral medulla
SPV: spinal trigeminal nucleus
TH: tyrosine hydroxylase
ts: solitary tract
VLM: ventrolateral medulla
vlPAG: ventral lateral periaqueductal gray
vmPFC: ventromedial prefrontal cortex
YFP: yellow fluorescent protein

